# The coordinated action of UFMylation and ribosome-associated quality control pathway clears arrested nascent chains at the endoplasmic reticulum

**DOI:** 10.1101/2025.01.17.633636

**Authors:** Milica Mihailovic, Aleksandra S Anisimova, Bu Erte, Ni Zhan, Ioanna Styliara, Yasin Dagdas, G Elif Karagöz

**Author notes:** equal contribution (Milica Mihailovic, Aleksandra S Anisimova).

## Abstract

Clearance of incomplete nascent polypeptides resulting from ribosomal stalling is essential for protein homeostasis. While ribosome-associated quality control (RQC) mechanisms that degrade these polypeptides are well-characterized in the cytosol, how stalled endoplasmic reticulum (ER)-bound ribosomes are cleared remains poorly understood. Stalled ER-bound ribosomes are marked by ubiquitin-fold modifier 1 (UFM1) on large ribosomal subunit protein RPL26, but the precise function and regulation of this process are unclear. Here, we demonstrate that canonical RQC factors associate with ribosomes stalled at the ER. Functional cellular assays using ER-targeted stalling reporters reveal that while ribosome splitting is a prerequisite for UFMylation of RPL26, the UFMylation persists without late RQC components that are involved in the clearance of arrested nascent chains (NEMF and LTN1). The UFM1 E3 ligase complex binds to and UFMylates the 60S-peptidyl-tRNA complex and, in concert with the canonical RQC pathway, facilitates the clearance of arrested polypeptides. Our findings reveal that UFMylation acts to maintain translational integrity at the ER.

## Introduction

To maintain cellular protein homeostasis, protein translation is constantly monitored by quality control factors. Distinct secondary mRNA structures, incompletely processed mRNAs, rare codons, or translation into poly(A) tail can lead to prolonged translational pausing and ribosomal stalling ^1–5^. Like a traffic jam, stalled ribosomes lead to ribosomal collisions and impaired translation ^6,7^. Also, incomplete nascent polypeptides are prone to aggregation, perturbing homeostasis ^2^. The highly conserved ribosome-associated quality control (RQC) pathway recognizes and recycles stalled ribosomes and degrades incomplete nascent chains to maintain proteostasis.

Cytosolic RQC mechanisms have been studied in depth (reviewed ^8–11^). Collided ribosomes are recognized by the E3 ligase ZNF598, which ubiquitinates 40S ribosomal proteins and halts translation ^5,12,13^. This is followed by the binding of the ribosome splitting factors ASC-1 complex/Pelo that recognize ubiquitinated stalled ribosomes ^14,15^ and split the leading ribosome to release the 60S-peptidyl-tRNA complex. After splitting, the 60S-peptidyl-tRNA complex is recognized by the NEMF/LTN1 complex. NEMF adds template-independent C-terminal alanine and threonine extensions (CAT-tails) to the stalled nascent chain, which facilitates the exposure of the nascent chains out of the ribosome tunnel for ubiquitination by the E3 ligase LTN1 ^16–20^. Subsequently, the nascent chain is released from the ribosome after tRNA cleavage by ANKZF1, extracted from the 60S ribosomal subunit by AAA ATPase VCP, and finally targeted for proteasomal degradation ^21–23^. The released 60S subunit is recycled and ready for translation initiation.

Translation into the endoplasmic reticulum (ER) membrane or the ER lumen poses topological and steric challenges for the RQC machinery to access the nascent chains on ER-bound ribosomes upon stalling. They also obstruct the translocon, hindering the synthesis and maturation of other proteins at the ER, which causes an additional burden on proteostasis ^24,25^. Our understanding of RQC for the ER-bound ribosomes that co-translationally translocate nascent chains into the ER is less understood ^26,27^. Recent studies showed that ribosomes stalled at the ER are marked by UFM1 (ubiquitin-fold modifier 1) at the large ribosomal subunit protein RPL26 ^28,29^. Similarly to ubiquitination, UFMylation proceeds through a cascade of enzymatic reactions catalized by an E1-activating enzyme (UBA5), an E2-conjugating enzyme (UFC1), and a complex E3-ligating enzyme (UFM1 E3 ligase complex) ^30–34^. The UFM1 E3 ligase complex consists of three components: UFL1, DDRGK1, and CDK5RAP3 (C53), and is tethered to the ER by the ER-membrane protein DDRGK1, providing specificity to the ER-bound ribosomes.

Accumulating evidence supports two, not necessarily mutually exclusive, models that explain the role of UFMylation in ER-localized translation. The first model suggests that ribosomal stalling at the ER induces RPL26 UFMylation, and UFMylation is involved in the clearance of stalled ribosomes and arrested polypeptides resulting from ribosomal stalling ^28,35^. The second model proposes UFMylation releases 60S ribosomal subunit from the translocon following the canonical termination of translation at the ER ^36,37^. Structural studies show preferential binding of UFL1 to empty 60S subunit and propose that simultaneous binding of UFL1 and tRNA is incompatible ^36,37^. However, this binding mode does not explain how the depletion of UFMylation machinery results in the accumulation of nascent arrested polypeptides in cells ^28,35^. To fill this gap in our knowledge, we dissected the interplay between the RQC and UFMylation machinery in clearing stalled ribosomes at the ER. We show that upon ribosomal stalling at the ER, UFL1 loads onto the 60S-peptidyl-tRNA complex. Through close collaboration with the RQC factors NEMF/LTN1, UFM1 E3 ligase complex facilitates clearance of the stalled peptides. Altogether, our findings underline the crucial role of the UFMylation-RQC crosstalk for clearing arrested incomplete polypeptides and ER-homeostasis.

## Results

### RQC factors associate with and act on UFMylated ribosomes

To discover regulators of UFMylated ribosomes, we purified them from mammalian cells and performed mass spectrometry (MS) analyses. To efficiently isolate UFMylated ribosomes, we generated an HCT116 cell line with endogenously tagged FLAG-UFM1 using the CRISPR-Cas9 gene editing approach. We then pelleted ribosomes by sucrose cushion sedimentation after inducing ribosomal stalling with anisomycin (ANS), which impairs peptidyl transferase activity of ribosomes ^38^. UFMylated ribosomes were subsequently enriched by FLAG immunoprecipitation (Fig. 1A). The MS analyses showed the expected enrichment of UFM1 E3 ligase complex components (UFL1, CDKRAP53, DDRGK1) in FLAG-immunoprecipitated samples from FLAG-UFM1 cells treated with anisomycin, compared to the input, and normalized to untreated cells as control (Fig. 1B, full list in Supplementary table 1). Additionally, we found that the RQC component NEMF is specifically associated with UFMylated ribosomes in a stalling-dependent manner (Figure 1B, C).

**Figure 1.**
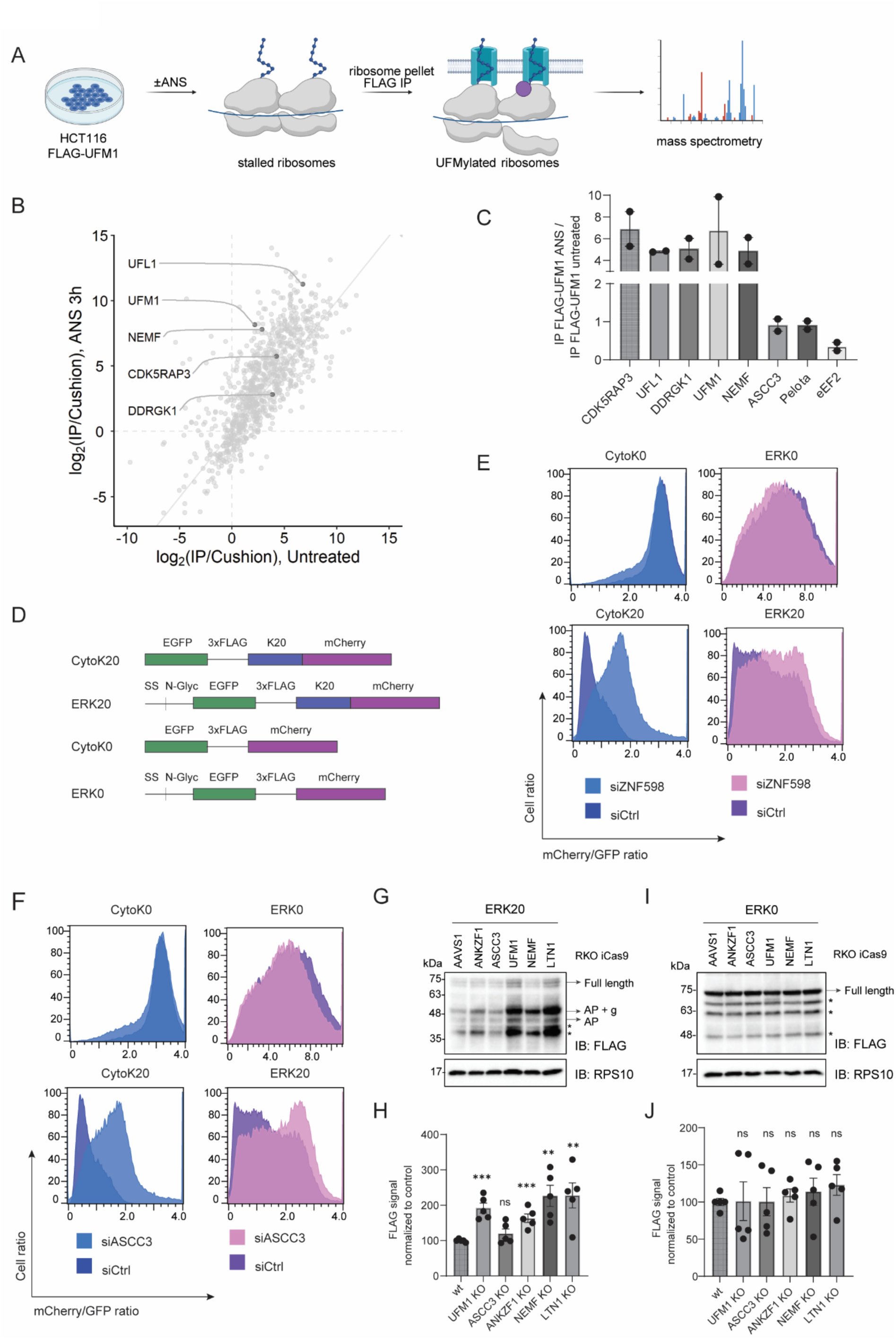
RQC and UFMylation machinery are needed to clear stalled peptides at the ER. (A) Scheme of mass spectrometry analysis of UFMylated ribosomes. (B) Scatter plot of proteins enriched with UFMylated ribosomes upon 3h 4μM ANS treatment compared to untreated HCT116 FLAG-UFM1 cells. (C) Enrichment of FLAG-eluates from ANS treated HCT116 FLAG-UFM1 cells over FLAG-eluates from untreated control HCT116 FLAG-UFM1 cells normalized to respective cushion samples, shown as log_2_ fold change, identified by mass spectrometry experiment from (A). (D) Schematic representation of control (K0) and stalling (K20) cytosolic and ER reporters. (E and F) Readthrough of reporters from (D) shown by mCherry/GFP ratio measured by FACS after 24h expression in HCT116 cells upon siRNA-mediated knockdown of ZNF598 (E) or ASCC3 (F) compared to non-targeting control siRNA. (G and I) Reporter accumulation shown by FLAG immunoblot in RKO iCas9 KO cell lysates upon 48h dox treatment to induce AAVS1 (non-targeting control), ANKZF1, ASCC3, UFM1, NEMF or LTN1 knockout, upon 24h ERK20 (G) or ERK0 expression (I) normalized to RPS10 loading control. AP – arrested peptide, g – glycosylation, * - degradation products. (H) Quantification of (G). (J) Quantification of (I).

The association of late RQC components with UFMylated ribosomes suggests that the cytosolic RQC machinery can also act on ribosomes stalled at the ER. To test whether the early RQC components recognize and split the ER-stalled ribosomes, we monitored the translational readthrough of well characterized stalling cytosolic or ER-targeted reporters ^28^ (Figure 1D) in a fluorescence-activated cell sorting (FACS) assay (Figure 1E, F). The ERK20 reporter consists of N-terminal signal sequence, N-glycosylation site, followed by EGFP and a poly-lysine stretch that mimics translation into polyA tail, and a C-terminal mCherry; while CytoK20 reporter has the same domain organization without the ER-targeting signal sequence and glycosylation site. To test whether early RQC machinery can access the ER-stalled ribosomes, we used siRNA to deplete ZNF598 and ASCC3, an ASC-1 complex component (Supplementary figure 1A), as depletion of ASCC3 leads to the 80% destabilization of the whole complex ^14^. Both the cytosolic (as shown earlier ^13^) and ER stalling reporters showed increased readthrough upon loss of ZNF598 and ASCC3 (Figure 1E-F). In contrast, control cytosolic CytoK0 and ER-targeted ERK0 reporters, which lack a stalling-inducing poly-lysine sequence, did not show a difference in readthrough upon depletion of the RQC components, as they don’t recruit the RQC machinery. Importantly, the loss of Pelo, which acts specifically on ribosomes stalled at the 3’-end of the RNAs ^24,39^, did not impact the readthrough of our internal ribosomal stalling reporters validating our findings (Supplementary figure 1B). Altogether, these results suggest that early RQC factors ZNF598 and ASCC3 can access and act on stalled ribosomes at the ER.

### RQC factors and UFMylation machinery collaborate to clear stalled peptides at the ER

After showing that early RQC components are essential for recognizing and splitting the ER-stalled ribosomes, we next tested the interplay between RQC factors and UFMylation machinery on the clearance of the model ER stalling substrate, ERK20. To deplete RQC factors, we used RKO cell lines that stably express doxycycline-inducible Cas9 (iCas9). Using lentiviral transduction, we introduced either specific guide RNAs (gRNAs) targeting the RQC components (ANKZF1, ASCC3, NEMF, LTN1) or the non-coding control locus AAVS1. In parallel, we used UFM1 iCas9 cells to impair UFMylation. Treatment of cells with doxycycline for 48 hours resulted in efficient depletion of the RQC factors or UFM1 (96% ANKZF1, 71% ASCC3, 75% NEMF and 69% LTN1 and 93% UFM1, Supplementary figure 1C). Importantly, we found that their depletion significantly stabilized the ER stalling reporter ERK20 (Figure 1G, H). Depletion of ASSC3 did not stabilize ERK20 reporter, since ASC-1 complex acts in early RQC. In contrast, as expected, depletion of these genes did not cause accumulation of the control ER reporter ERK0 (Figure 1I, J).

We next tested the downstream degradation pathway of cytosolic and ER stalling reporters through proteasomal inhibition with MG132 and autophagy inhibition with bafilomycin treatment. Both reporters are stabilized by the inhibition of proteasome and autophagy machineries, yet the proteasomal inhibition showed a higher accumulation of the reporters indicating its major contribution to their clearance (Supplementary figure 1D). Taken together, we show that both the UFMylation and the RQC machinery are involved in clearing arrested nascent peptides on stalled ribosomes at the ER.

### Recognition and splitting of the ER-stalled ribosomes precedes UFMylation

Next, we investigated the crosstalk between the RQC pathway and UFMylation. To reveal the order of events and the interdependence of these pathways, we first assessed whether the loss of RQC components impacts UFMylation of the ER-stalled ribosomes. To this end, we used RKO iCas9 cell lines expressing gRNAs against RQC components (Supplementary figure 1C). Under steady-state conditions, depletion of ZNF598, ASCC3, or PELO did not impact UFMylation levels (Figure 2A, lanes 1-4, Figure 2B). However, upon ANS treatment, we noticed a significant decrease in UFMylation levels upon depletion of ZNF598, ASCC3, or Pelo compared to control (AAVS1) (Figure 2C, lanes 1-3, Figure 2D). These results were corroborated in HCT116 cell line using siRNA depletion of RQC components (Supplementary figure 2A, B).This data indicated that ribosome splitting is required for UFMylation of ribosomes at RPL26. Additionally, we detected ZNF598-dependent ubiquitination of 40S ribosome protein RPS10 on disomes by polysome profiling in HCT116 cells treated with ANS, while UFMylation was mainly enriched in 60S fractions, suggesting that early RQC-mediated ubiquitination and UFMylation have different substrates (Supplementary figure 2C).

**Figure 2.**
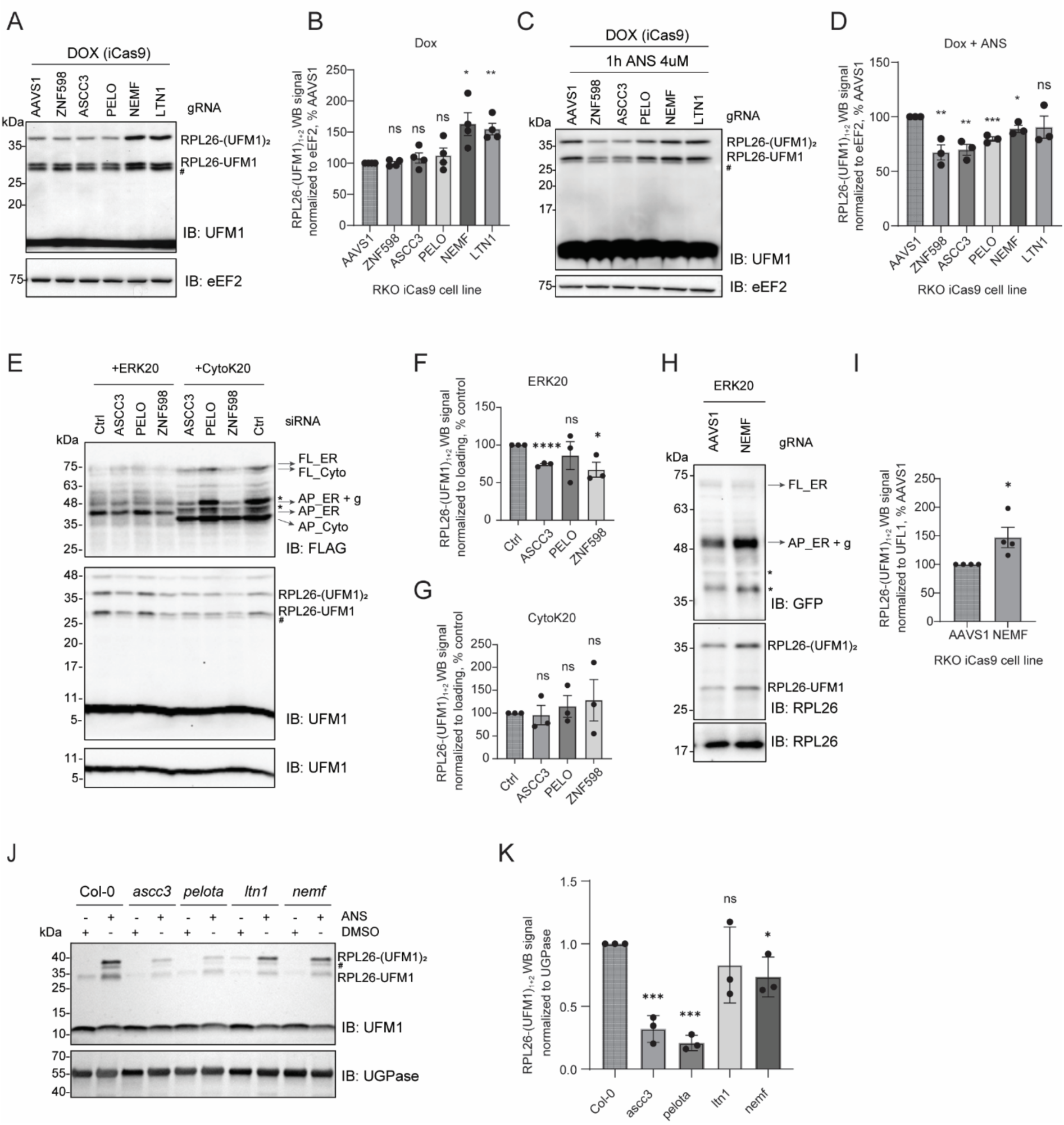
RPL26 is UFMylated on post-split 60S subunit upon ribosomal stalling at the ER. (A and C) UFMylation levels visualized by UFM1 immunoblot in RKO iCas9 cells upon 48h dox treatment to induce RQC components or non-targeting AAVS1 knockout in untreated (A) or 1h 4μM ANS treated cells (C). (B) Quantification of (A). (D) Quantification of (C). (E) UFMylation levels visualized by UFM1 immunoblot in HCT116 cells upon 72h siRNA-mediated knockdown of RQC components or non-targeting control, and 24h expression of ERK20 or CytoK20. (F and G) Quantification of (E) for cells expressing ERK20 (F) or CytoK20 (G) shown by FLAG signal normalized to loading control. (H) UFMylation levels visualized by RPL26 immunoblot in RKO iCas9 cells upon 48h dox treatment to induce non-targeting AAVS1 or NEMF knockout upon 24h expression of ERK20 reporter. (I) Quantification of (H). (J) 7-day-old *Arabidopsis* seedlings of wildtype (Col-0) and RQC component mutant (*ascc3*, *pelota*, *ltn1*, *nemf*, *rqc1*, *edf1*) were treated with either DMSO or 100 μM ANS for 16 hours. The UFMylation level was tested via immunoblotting using anti-UFM1 antibody, with UGPase was introduced as loading control. The data shown are representative of three biological replicates. (K) Quantification of (J) for ANS-treated samples. FL – full length, AP – arrested peptide, g – glycosylation, # - UFC1-UFM1 complex ^30^, * - degradation products.

Notably, the depletion of RQC factors NEMF and LTN1, which facilitate ubiquitination and clearance of the nascent chain, increased RPL26 UFMylation levels under steady-state conditions (Figure 2A, lanes 5,6, Figure 2B), suggesting that RQC continuously surveils aberrant translation. We anticipate that depletion of NEMF and LTN1 impairs the clearance of the stalled ribosomes at the ER (as we showed in Figure 1G, H), leading to the accumulation of the UFMylated 60S-peptidyl-tRNA complex intermediates. Surprisingly, NEMF and LTN1 depletion did not largely impact RPL26 UFMylation upon ANS treatment (Figure 2C, lanes 5,6, Figure 2D). Compared to steady-state conditions in Figure 2A, this difference could be due to cell-wide ribosomal stalling induced by ANS treatment exhausting the UFMylation machinery. These data suggest that the UFMylation machinery acts at capacity, therefore, the depletion of NEMF and LTN1 during ANS treatment does not lead to a further increase.

To uncover ER-specific cross-talk between the RQC and UFMylation machinery, we induced ribosomal stalling in a more specific manner by expressing cytosolic or ER stalling reporters ERK20 and CytoK20 in HCT116 cells. As shown before ^28^, while ERK20 expression increased RPL26 UFMylation, CytoK20 did not (Figure 2E). The siRNA-mediated knockdowns of ZNF598, and ASCC3 resulted in a significant decrease in RPL26 UFMylation levels in cells expressing ERK20, compared to control siRNA, while PELO knockdown did not impact UFMylation (Figure 2E, lanes 1-4, Figure 2F). In contrast, cells expressing CytoK20 showed no difference in RPL26 UFMylation levels upon knockdowns of early RQC components ZNF598, and ASCC3 (Figure 2E, lanes 5-8, Figure 2G). Similarly, their depletion did not impact UFMylation levels in cells expressing control ERK0 reporter (Supplementary figure 2D, E). Therefore, the depletion of ZNF598 and ASC-1 complex decreases UFMylation, specifically in cells expressing ER stalling reporter ERK20. We also noticed an increase in UFMylation upon NEMF depletion in RKO iCas9 cells expressing ERK20 reporter (Figure 2H, I) showing that UFMylation of RPL26 is not dependent on NEMF.

Given the conservation of the UFMylation system across eukaryotes, we proceeded to investigate the evolutionary conservation of this mechanism in plants. To this end, we evaluated the effect of RQC machinery on RPL26 UFMylation using *Arabidopsis thaliana* mutant lines. UFMylation levels were assessed upon ANS treatment in *ascc3*, *pelota*, *ltn1*, and *nemf* mutant lines compared to wt (Columbia Col-0 ecotype). Knocking out *ascc3* and *pelota* significantly decreased UFMylation levels upon stalling induction by ANS treatment compared to the wild-type (Figure 2J, K). Additionally, we noticed a slight decrease in nemf mutant line (Figure 2J, K). These results confirm our hypothesis that the order of events upon ribosome stalling at the ER is conserved across eukaryotes.

To sum up, we show that upon stalling at the ER, ribosomes are recognized by ZNF598 and split by ASC-1 complex. The ribosome splitting precedes and is required for UFMylation of the ER-bound ribosomes upon stalling. At the same time, the depletion of the late RQC components NEMF and LTN1, which are involved in the clearance of the arrested peptides on the 60S, does not impair the UFMylation of RPL26. These results indicate that the nascent polypeptide release is not necessary for the UFM1 E3 ligase complex to access and UFMylate the 60S. Taken together, we mechanistically show that translational stalling at the ER induces UFMylation of post-splitting 60S-peptidyl-tRNA complexes, and this mechanism is evolutionarily conserved in plants and mammals.

### UFM1 E3 ligase complex binds nascent chain-associated 60S-peptidyl-tRNA complex at the ER

The cryo-electron microscopy structures of UFL1 complex bound to 60S ribosomes revealed that UFM1 E3 ligase complex forms extensive contacts with the 60S in a clamp-like architecture extending from tRNA-binding sites to the peptide exit tunnel with all the three subunits of complex contributing to this interaction ^36,37^. Based on these structures, it was proposed that UFL1 binding blocks the tRNA-binding site and these two binding events are mutually exclusive. However, those structures cannot explain how the loss of the components of the UFM1 E3 ligase leads to the accumulation of ER stalling reporters. Namely, it is not clear how machinery that only works on post-termination ribosomes could result in the accumulation of the arrested nascent polypeptides (Fig. 1G, H). To this end, we next dissected whether binding of the UFM1 E3 ligase complex occurs before or after nascent chain release from ER-bound 60S subunits.

Depletion of the RQC factors involved in the nascent chain clearance increased UFMylation of RPL26, suggesting that the UFM1 E3 ligase complex can access and UFMylate nascent chain-associated 60S subunits (Figure 2A, B, H, I). The catalytic component of the UFM1 E3 ligase complex, UFL1, forms the central scaffold of the UFM1 E3 ligase complex and forms extensive contacts with both other components of the E3 ligase complex as well as the 60S. The structural model of UFL1 displays a short N-terminal α-helix followed by one partial winged-helix (pWH), five WH motifs, a bipartite coiled-coil (CC) domain with a disordered region and a C-terminal globular domain (CTD), which contacts the 28S ribosomal RNA (rRNA) occluding all three tRNA-binding sites (Figure 3A, B). Based on our data and data from others ^36,37^, we hypothesized that the extensive contacts formed between the UFM1 E3 ligase complex and the 60S would allow for sufficient affinity for this interaction to occur even though the tRNA site is occupied. Notably, the structural models of the UFL1 show that CTD displays high degree of conformational freedom due it being connected to the rest of the protein with a disordered segment (Figure 3A). Overlaying the published structures of 60S ribosomes with UFL1 complex and NEMF/LTN1 shows a possibility of a hybrid state with a possible rearrangement of the CTD of UFL1 and NEMF/LTN1 at the P-site (Figure 3B). Supporting our data, deletion of the CTD of UFL1 does not entirely abolish UFMylation of the RPL26 in cells^36^. Therefore, we speculated that UFL1 could undergo conformational rearrangements to enable its binding to 60S-peptidyl-tRNA complex (Figure 3B).

**Figure 3.**
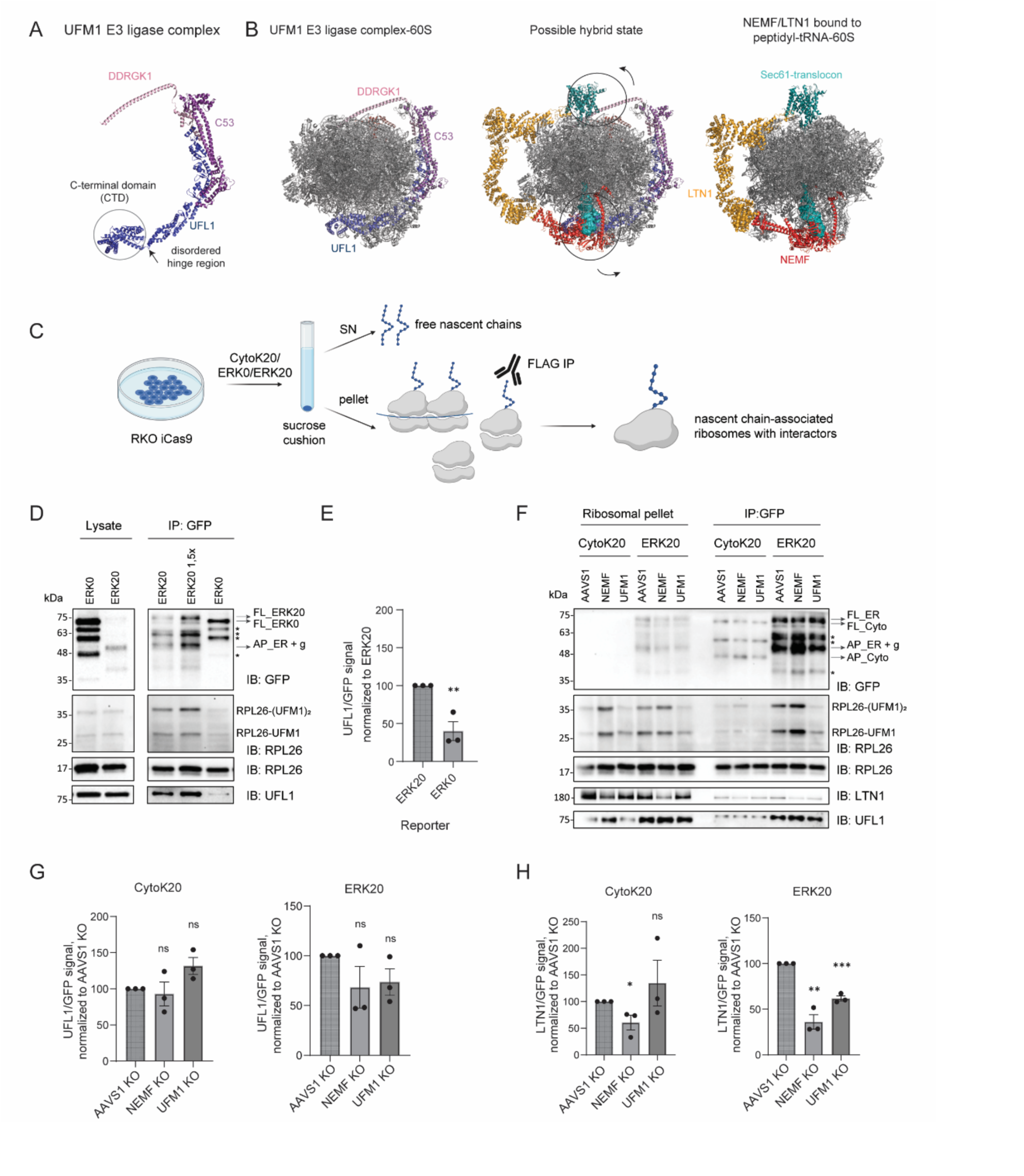
UFL1 UFMylates nascent chain-bound 60S subunits. (A) The cryostructural model of UFM1 E3 ligase complexes shows flexibility around the CTD region of UFL1 (pbd: 8ohd). (B) Left: The structure of UFM1 E3 ligase complex bound to 60S subunit (pbd: 8ohd), right: the structure of NEMF/LTN1 complex bound to peptidyl-tRNA-60S complex (pdb: 3j92, 8agw) docked on translocon (pdb:3j7r), middle: overlay of two structures shows a possible hybrid state with conformational rearrangements of UFL1 and DDRGK1. (C) Experimental approach for detecting interactors of nascent chain-associated ribosomes. (D) Ribosomal pellet was obtained by sucrose cushion centrifugation followed by GFP immunoprecipitation from RKO iCas9 parental cell line upon 24h expression of ERK0 or ERK20 reporter. Protein levels are analyzed by immunoblotting. (E) Quantification of UFL1 association normalized to eluate nascent chain levels from (D). (F) Ribosomal pellet was obtained by sucrose cushion centrifugation followed by GFP immunoprecipitation from RKO iCas9 AAVS1 (non-targeting control), NEMF or UFM1 KO cell line upon 24h expression of CytoK20 or ERK20 reporter. Protein levels are analyzed by immunoblotting. (G) Quantification of UFL1 association normalized to eluate nascent chain levels from (F) for cells expressing CytoK20 or ERK20. (H) Quantification of LTN1 association normalized to eluate nascent chain levels from (F). FL – full length, AP – arrested peptide, # - UFC1-UFM1 complex ^30^, * - degradation products.

To experimentally test this model, we assessed whether the UFL1 complex associates with the 60S-peptidyl-tRNA complex following ribosome splitting. To this end, we enriched nascent chain-bound ribosomes by immunoprecipitating ER stalling reporter ERK20 or control ER reporter ERK0 in RKO cells (Figure 3C). To enrich for the ribosome-associated nascent chains, we first isolated ribosomes via sucrose cushions and performed immunoprecipitation (Figure 3C). We found that UFL1 specifically associates with ERK20 on the nascent chain-bound ribosomes, while we did not observe UFL1 binding to ERK0 (Figure 3D). Quantification of three replicates showed a significant difference in binding between ERK0 and ERK20, showing that UFL1 binding depends on ribosome stalling at the ER (Figure 3E). Importantly, we also observed an enrichment of the UFMylated RPL26 in the eluates from ERK20-expressing cells (Figure 3D, lanes 3-5). These data indicated that UFL1 can bind and actively UFMylate 60S-peptidyl-tRNA complex with the arrested nascent chains at the ER.

To test whether the association of NEMF/LTN1 with the 60S is required for UFL1 binding to the 60S-peptidyl-tRNA complex at the ER, we conducted similar experiments upon NEMF depletion in RKO iCas9 cells expressing CytoK20 or ERK20 reporters (Figure 3C). IP analyses showed specific interaction of UFL1 with the ERK20-stalled ribosomes while it did not associate with the ribosomes stalled in the cytosol (CytoK20) (Figure 3F, G). Moreover, UFL1 binding was unaffected by NEMF depletion, supporting the evidence that UFMylation precedes the activity of the late RQC components (Figure 3G, right panel). Notably, upon NEMF depletion, EK20 pulldowns displayed higher levels of UFMylated RPL26, further supporting our model that UFL1 binds to and UFMylates 60S-peptidyl-tRNA complex before nascent chain release (Figure 3F, lanes 10,11). The UFM1 depletion did not impair association of UFL1 with ERK20 expressing ribosomes, indicating that this step precedes stabilization of UFL1-ribosome complexes via UFM1 attachment (Figure 3F, lanes 10, 12). We also noticed a NEMF-dependent association of LTN1 to the 60S-peptidyl-tRNA complexes both for cytosolic and ER-stalled ribosomes (Figure 3H), in accordance with previous data ^40^. Interestingly, we show that LTN1 binding decreases upon UFM1 KO for ER-stalled ribosomes (Figure 3H). This can be explained by destabilization of the translocon-60S association by UFMylation of the 60S-peptidyl-tRNA complexes, thus allowing better access of LTN1 to the nascent chain (Figure 3B). In summary, we demonstrate that the UFM1 E3 ligase complex binds to 60S-peptidyl-tRNA on the ER following ribosomal stalling. This interaction does not depend on binding of the NEMF/LTN1 complex and facilitates the clearance of arrested polypeptides on ER stalled ribosomes.

## Discussion

The best-described substrate of the UFMylation machinery is the large ribosomal subunit protein RPL26, initially discovered to be increasingly UFMylated upon ribosomal stalling at the ER ^28,29^. However, the primary function of this event has remained poorly understood. Recent cryo-EM structures uncovered a novel function of the UFMylation machinery in recycling the 60S subunits after translational termination at the ER by releasing them from the translocon ^36,37^. These structures also showed that UFL1 occupies the tRNA binding sites at the 60S, suggesting that the UFM1 E3 ligase binds to 60S subsequent to translation termination ^36,37^. Therefore, the role of UFMylation in clearing ER-stalled ribosomes remained unclear.

Here, using IP-MS analyses, we found that RQC components are associated with the UFMylated ribosomes (Figure 1B, C). To confirm the role of RQC factors on clearing ER stalled ribosomes, we used genetic depletion of RQC machinery and found that ribosomes stalled upon ERK20 expression recruit the RQC E3 ligase ZNF598 and the ribosome splitting factor ASC-1 complex (Figure 1E, F, Figure 2E-I). Similar to RQC events upon cytosolic stalling, the stalled ribosomes at the ER required the activity of ANKZF1, NEMF and LTN1 for clearance of arrested polypeptides since the loss of these components stabilized the ER stalling reporter (Figure 1G, H). Likewise, impaired UFMylation specifically stabilized ERK20 substrate in line with the published work (Figure 1G, H) ^28^, indicating that both the RQC and the UFMylation machinery are involved in the clearance of the stalled ribosomes at the ER.

After demonstrating the role of the RQC and the UFMylation machinery in the clearance of the ER-stalled ribosomes, we assessed the sequence of events and interdependence of these pathways by testing the impact of the loss of RQC factors on RPL26 UFMylation upon ribosome stalling. The RPL26 UFMylation levels decreased upon knockdown or knockout of ZNF598 or ASCC3 in cells treated with anisomycin, a drug inducing ribosomal stalling (Figure 2 C, D, Supplementary figure 2A, B) as well as upon expression of the specific ER stalling reporter ERK20 (Figure 2E-I). Experiments performed in plants (*Arabidopsis thalian*a) validated those findings (Figure 2J, K), concluding that UFMylation happens on post-split 60S ribosomes upon ribosomal stalling at the ER and that this mechanism is conserved from plants to mammals.

Previous structural data showed that UFL1 mainly binds to the 60S ribosomal subunit in a way that excludes the binding of the 40S subunit, translocon, or tRNA ^36,37^. These data cannot be reconciled with the data showing increased UFMylation of RPL26 upon ribosome stalling. We showed that UFL1 associates with ribosomes expressing ER stalling reporter ERK20, but not the cytosolic stalling reporter CytoK20 or the ER-targeted control reporter ERK0 (Figure 3D, E). Immunoprecipitation of the ribosome-attached ERK20 stalling reporter showed that these ribosomes were already UFMylated at RPL26, demonstrating that UFM1 E3 ligase can act on nascent chain-loaded ribosomes, contradicting the previous model (Figure 3D, E) ^36,37^. Importantly, the depletion of NEMF and UFM1 did not impact UFL1 binding to the nascent chain-containing ribosomes (Figure 3F, G). Moreover, we readily detected UFMylated RPL26 in the nascent chain immunoprecipitates from NEMF KO cells, showing that UFMylation happens independently of NEMF binding (Figure 3F). Supporting our findings, single particle cryo-electron microscopy studies of the native UFM1 E3 ligase complexes isolated from cells showed a small population of 60S with a weak extra density in the peptide exit tunnel, which could possibly represent a nascent polypeptide chain ^36^. To sum up, we found that the 60S-peptidyl-tRNA complex formed upon ribosome splitting can serve as a substrate for UFM1 E3 ligase complex and that the NEMF/LTN1-dependent release of the nascent chain is not necessary for RPL26 UFMylation.

In cytosolic ribosomes, LTN1 binds close to the ribosome exit tunnel in the vicinity of the stalled nascent polypeptides emerging from the ribosome ^40^. In the ER-bound ribosomes exit tunnel is occupied by the translocon ^41,42^. The RQC pathway was proposed to act on ER-stalled ribosome through exposure of the nascent polypeptide in the cytosol at the ribosome-translocon contact site ^43^. As depletion of the UFMylation machinery stabilizes the arrested nascent chains, and the binding of LTN1 to 60S at the ER is enhanced by UFMylation (Figure 3F, H), we speculate that, similar to what has been proposed for the post-termination empty 60S subunits, binding of the UFM1 E3 ligase complex destabilizes the 60S-translocon interactions and allows the E3 ligase LTN1 to access the arrested nascent chain for ubiquitination and subsequent degradation (Figure 3B).

Both ubiquitin proteosome system and lysosomal degradation was proposed be involved in clearance of stalled nascent peptides at the ER ^28,33,35^. Notably, the previous work on the ERK20 reporter used here showed that it is mainly degraded by lysosomes and neither proteosome inhibition nor NEMF depletion stabilized the reporter ^28^. However, we show a clear stabilization of the same reporter preferentially by proteosome inhibition (Supplementary figure 1D), and NEMF depletion (Figure 1G, H). Supporting our results, an alternative ER stalling reporter containing a folded VHP domain and a poly-lysine stretch, was degraded primarily by the proteasome, and showed a NEMF-dependent stabilization ^35^. We therefore conclude that while RQC and ubiquitin proteosome system play the major role to clear stalled ribosomes, depending on the cell type and the expression level, the arrested polypeptides at the ER can be cleared by complementary degradation pathways including ER-phagy and lysosomal pathways ^28,33^.

Altogether, our data converge on the following model: stalling of the ER-bound ribosomes recruits canonical ribosome-associated quality control machinery. First, the E3 ligase ZNF598 recognizes the collided ribosomes and stabilizes them by ubiquitinating small subunit proteins ^13^. This is followed by the binding of the ASC-1 complex that splits the leading ribosome, leaving a 60S-peptidyl-tRNA complex (Figure 4). The 60S-peptidyl-tRNA complex is then recognized and UFMylated by UFM1 E3 ligase independently of downstream RQC components’ activities. RPL26 UFMylation allows better access of LTN1 to the nascent chain that gets ubiquitinated and targeted for downstream proteasomal degradation to restore cellular proteostasis. This process shows high evolutionary conservation in plants and mammals and highlights the importance of fine regulation and complementarity of ribosomal quality control processes.

**Figure 4.**
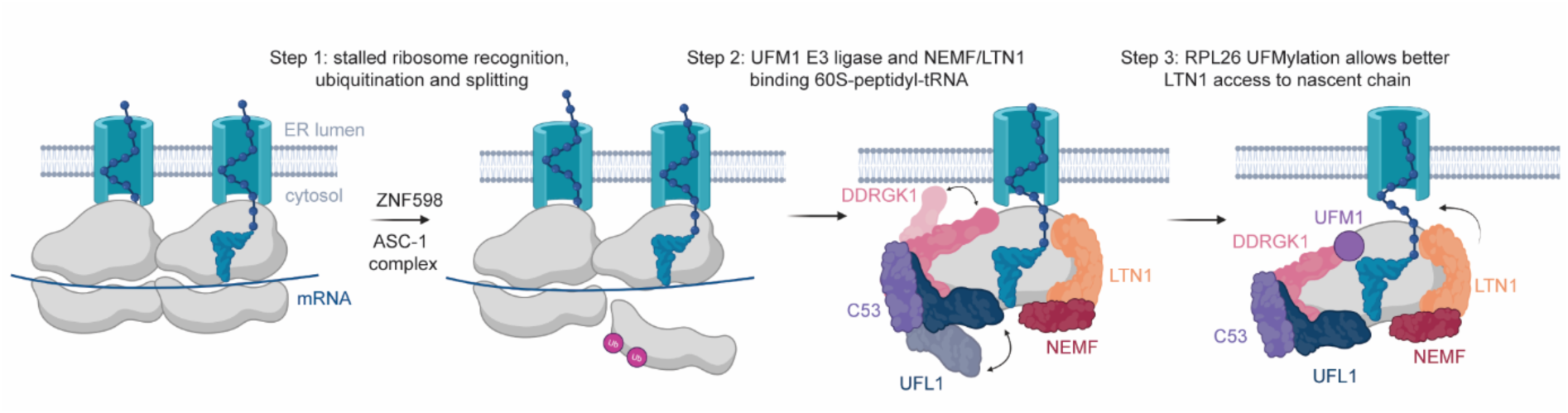
UFMylation machinery acts together with RQC to clear arrested peptides at the ER. Upon ribosomal stalling at the ER, ZNF598 recognizes and ubiquitinates 40S proteins, followed by ASC-1 complex-mediated splitting of the leading ribosome. The remaining 60S-peptidyl-tRNA complex is recognized and UFMylated by the UFM1 E3 ligase complex, independently of downstream RQC factors. UFMylation allows easier access to nascent chain that gets ubiquitinated by LTN1 with the help of NEMF. The nascent chain gets degraded by the proteasome and the 60S subunit is recycled.

## Acknowledgements

We thank Kitti Csalyi and Thomas Sauer at Max Perutz Labs Biooptics FACS facility and Elisabeth Roitinger at Vienna BioCenter Proteomics Core Facility for their help. We are thankful to Gijs Versteeg (Max Perutz Labs, Vienna) and Johannes Zuber (IMP, Vienna) for the help with the iCas9 RKO cel llines and lentiviral transduction, and to Yihong Ye (NIH, Bethesda) for CytoK0, CytoK20, ERK0, and ERK20 plasmids. This research was funded in whole or in part by the Austrian Science Fund (FWF) [FWF-W1261, FWF-DOC 177B] to GEK, the [FWF-SFB F79] and Vienna Science and Technology Fund, WWTF-LS21 Chemical Biology to GEK and YD and European Research Council Grant (Project number: 101043370) to YD. ASA and MM are supported by the DOC fellowship of Austrian Academy of Sciences. For open access purposes, the author has applied a CC BY public copyright license to any author-accepted manuscript version arising from this submission. Schemes are generated in BioRender.

## Figure Legends

**Supplementary Figure 1.**
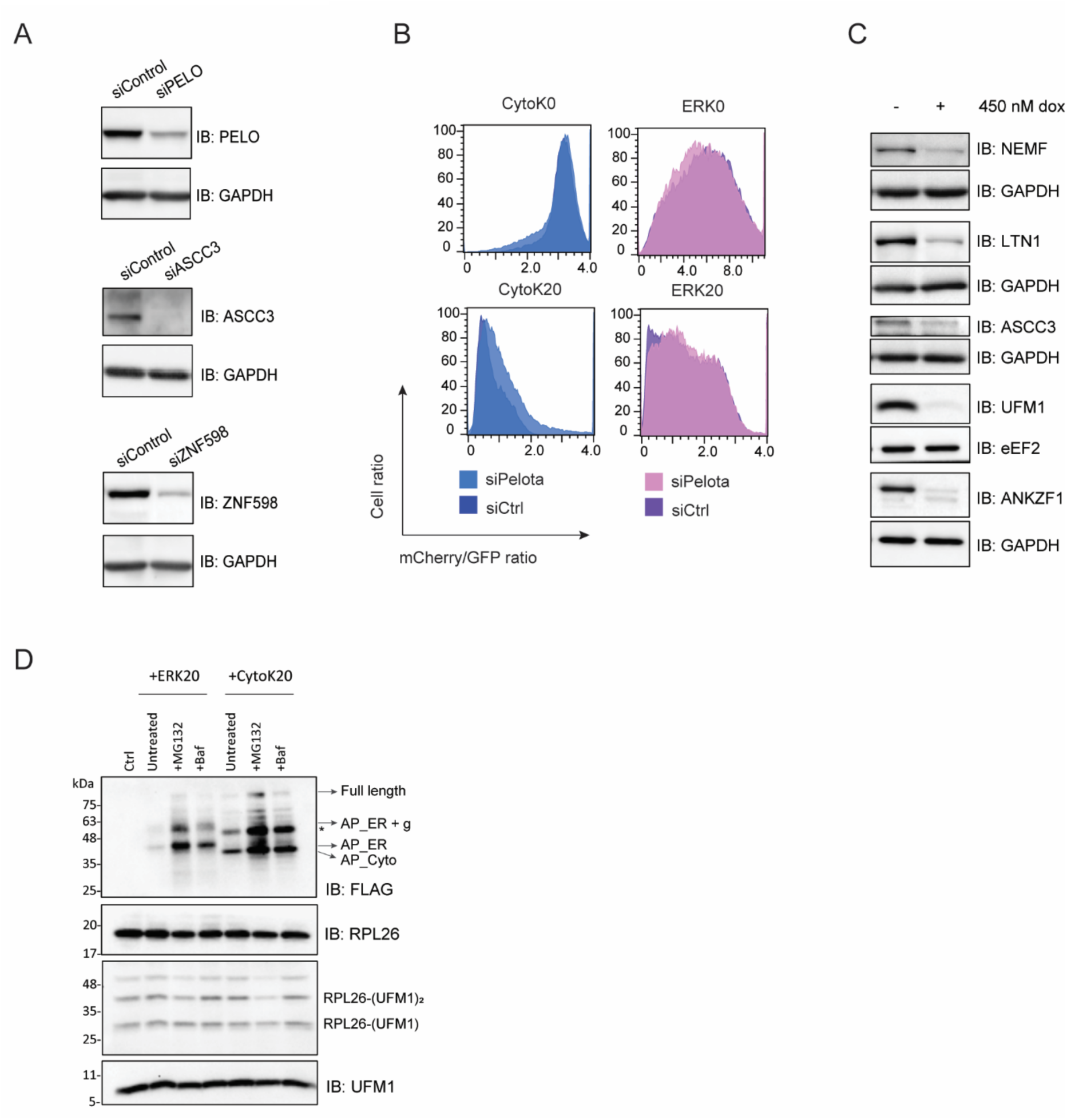
(A) Scatter plot of proteins enriched with UFMylated ribosomes upon 3h 4μM ANS treatment compared to untreated HCT116 FLAG-UFM1 cells, shown as log_2_ fold change of eluates over ribosome cushions (second replicate from Figure 1B). (B) Immunoblots showing knockdowns of RQC proteins Pelota, ASCC3, or ZNF598 upon 72h siRNA treatment in HCT116 cells. GAPDH is used as a loading control. (C) Readthrough of reporters from Fig. 1D shown by mCherry/GFP ratio measured by FACS after 24h expression in HCT116 cells upon siRNA-mediated knockdown of Pelota compared to non-targeting control siRNA. (D) Immunoblots showing knockouts of RQC proteins NEMF, LTN1, ASCC3, ANKZF1, or UFM1 knockout upon 48h dox treatment in RKO cells. GAPDH or eEF2 are used as a loading control. (E) Immunoblots showing reporter accumulation upon 24h expression of CytoK20 or ERK20 in HCT116 cells after treatment with either 20 nM MG132 for 3h, 10 nM Baf for 16h, or untreated as a control. AP – arrested peptide, * - degradation products.

**Supplementary Figure 2.**
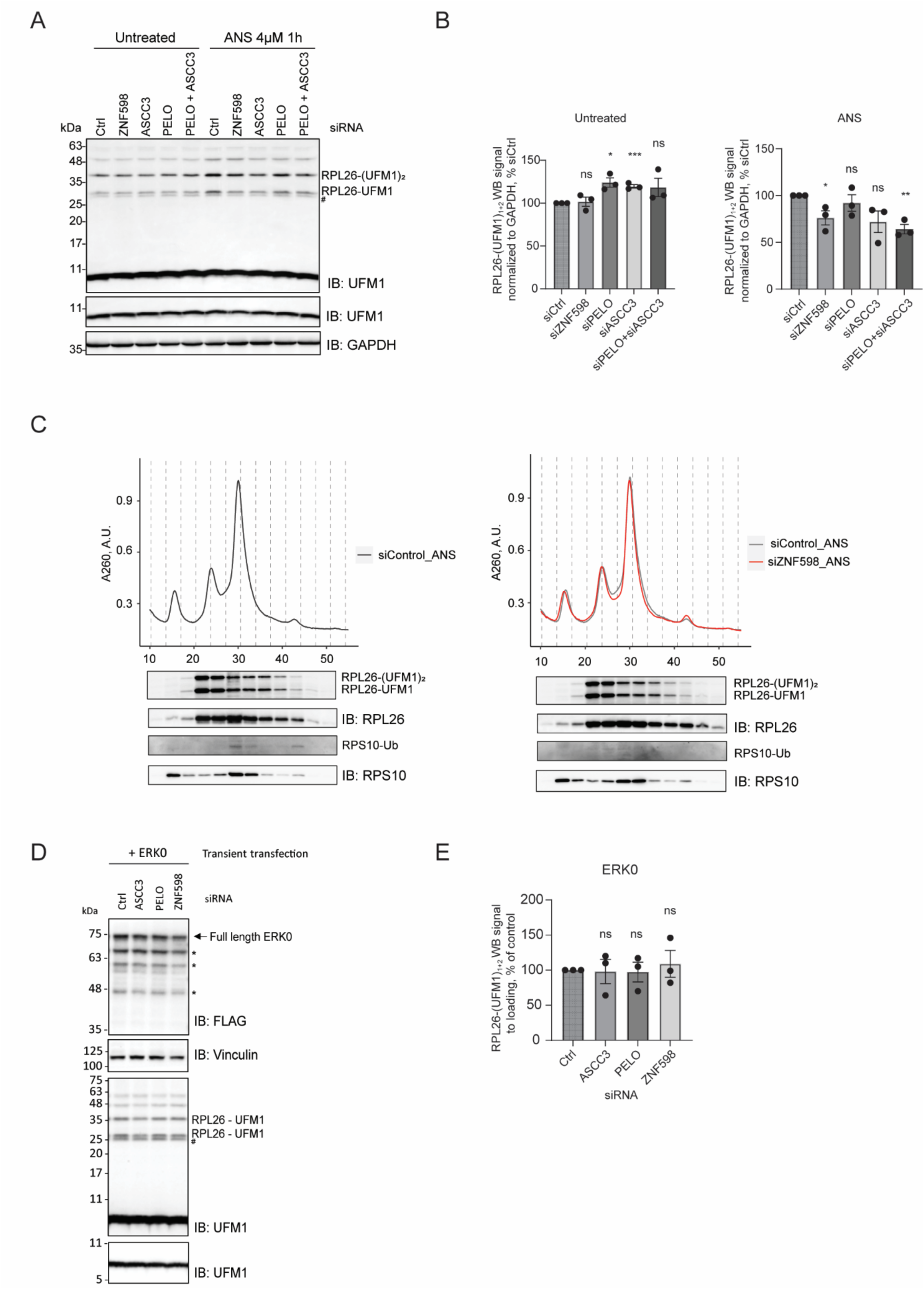
(A) UFMylation levels visualized by UFM1 immunoblot in untreated or 1h 4μM ANS treated HCT116 cells upon 72h siRNA-mediated knockdown of RQC components or non-targeting control. (B) Quantification of (A). (C) Top: polysome profiles from HCT116 cells upon 72h knockdown of ZNF598 (right panel) or control siRNA (left panel). Bottom: immunoblots showing RPL26 and RPS10 protein levels from corresponding fractions.(D) UFMylation levels visualized by UFM1 immunoblot in HCT116 cells upon 72h siRNA-mediated knockdown of RQC components or non-targeting control, and 24h expression of ERK0. (E) Quantification of (D). # - UFC1-UFM1 complex, * - degradation products.

## Materials and Methods

### Mammalian cell culture

HCT116 tetON (doxycycline-inducible) OsTIR1 cells obtained the Masato Kanemaki lab ^44^ were cultured in McCoy’s 5A (modified) medium (Sigma, M9309) supplemented with 10% fetal bovine serum (Gibco, 10437028), 2 mM L-Glutamine (Sigmam, G7513), 1% Pen/Step (Sigma, P0781). RKO-Dox-Cas9 cells (expressing doxycycline-inducible Cas9), a kind gift from from Johannes Zuber lab ^45^ were cultured in RPMI-1640 media (Sigma, R8758) supplemented with 10% fetal bovine serum (Gibco, 10437028), 2 mM L-Glutamine (Sigmam, G7513), 1% Pen/Step (Sigma, P0781), 1x non-essential amino acids (Thermo Scientific, 11140050), and 1 mM sodium pyruvate (Gibco, 11360070). Cell lines were grown in a humidified incubator at 37°C and 5% CO_2_. All cell lines were regularly tested for *Mycoplasma* infection with the EZ-PCR™ Mycoplasma Detection Kit (Biological Industries).

### Mammalian cell line generation

RKO-Dox-Cas9 (iCas9) cell lines for doxycycline inducible knockout of the genes encoding RQC components were established using lentiviral transduction with lentiviral particles (produced as described in ^46^ containing Dual-sgRNA_hU6-mU6 vectors described in ^45^ expressing two sgRNAs (Supplementary table 2) from human and mouse U6 promoters and eBFP2 from a PGK promoter. The eBFP2-positive cells were FACS sorted at BD FACSMelody™ Cell Sorter at Max Perutz Labs BioOptics FACS Facility.

To endogenously tag UFM1 with N-terminal 3xFLAG sequence human UFM1 N-terminal homology arms sequence was amplified using HindIII_UFM1_N_HAs_F and XhoI_UFM1_N_HAs_R primers. The UFM1 homology arms were inserted into the pMK344 plasmid backbone (Addgene #121179) using HindIII and XhoI restriction sites. To remove the BamHI site the plasmid was treated with HindIII and XbaI restriction enzymes and ligated through annealed overlapping oligos (pMK344_oligo_HindIII_XbaI_F/R). The BamHI and SalI restriction sites were introduced to the homology arms using BamHI_UFM1_N-HA and SalI_UFM1_N-HA primers. The BSD-P2A-3xFLAG sequence was cloned using the pMK347 (BSD-P2A-mAID) plasmid (Addgene #121181) which was digested with the Esp3I and BamHI enzymes to remove P2A-mAID and ligated using P2A-3xFLAG template made from annealed oligos (3xFLAG_pMK347_F/R) with cohesive ends. The BSD-P2A-3xFLAG sequence was inserted into the plasmid with N-terminal homology arms of UFM1 using SalI and BamI restriction sites resulting in the HDR template. The PAM sites were mutated in the HDR template using site-directed mutagenesis with a primer pair UFM1_gRNA_517_PAM_mut_F/R. To clone the Hygro-P2A-3xFLAG HDR template Hygro-P2A-3xFLAG was amplified from pMK344 with KS_F and P2A_3xFLAG_BamHI_R primers and cloned to replace the BSD-P2A-3xFLAG in the final UFM1 HDR template with mutated PAM sites. All primer sequences are listed in Supplementary table 2.

To introduce the 3xFLAG tag to the N-terminus of the endogenous UFM1 HCT116 tetON OsTIR1 cells ^44^ were transiently transfected with 1:1:2 ratio mixture of the BSD-P2A-3xFLAG HDR, Hygro-P2A-3xFLAG HDR template plasmids and pSpCas9 (BB)-2A-GFP (PX458) (plasmid #48138, Addgene) ^47^ targeting the first exon of the UFM1 gene (Supplementary table 2) using the Fugene HD (Promega) reagent according to the manufacturer’s instructions. 24 hours after transfection cell were collected by trypsinization and plated in 1:200 dilution in standard culture media (McCoy’s 5A (modified) with 10% FBS, 2 mM L-Glutamine, 1% Pen/Step). On the next day the media was supplemented with 100 µg/mL of Hygromycin B Gold (InvivoGen, #ant-hg) and 10 µg/mL of Blasticidin S Hydrochloride (InvivoGen, #ant-bl-05). Cells were grown in selection media until visible colonies were formed and expanded in 96-well plates. The homozygous insertion was verified using western blotting with anti-UFM1 (ab108062, Abcam) and anti-GAPDH (10494-1-AP, Proteintech) antibodies and genotyped using DirectPCR Lysis-Reagent Cell (Peqlab, VWR) with hsUFM1_HAs_F, hsUFM1_HAs_R, HygR_F, and BSDR_F primers.

### Co-Immunoprecipitation mass spectrometry of 3xFLAG-UFM1

For mass spectrometry of 3xFLAG-UFM1 co-immunoprecipitation (co-IP) ten 15 cm (diameter) dishes of 80% confluent 3xFLAG-UFM1 HCT116 per condition were used. Cell culture media was exchanged to fresh one 16 hours prior collection. To induce ribosome stalling cells were treated with 4 μM anisomycin for 3 h. Cells were washed with warm (37°C) PBS supplemented with 100 μg/mL cycloheximide, lysed on the plate with ice-cold lysis buffer (20 mM HEPES pH 7.3, 150 mM KCl, 5 mM MgCl_2_, 1% Triton X-100, 100 mg/mL cycloheximide, 1 mM DTT, 1x EDTA-free protease inhibitor cocktail), scraped, and transferred to 2 mL RNase-free tubes. Cell lysates were further incubated on ice for 10 min with intermittent vortexing, passed three times through the 27G needle, and clarified on a table-top centrifuge at maximum speed (20,000 g) for 20 min at 4°C. Total RNA concentration in the lysate was estimated using OD A260 measurement on Nanodrop. The lysate concentrations were equalized and brought to 1.5 mL with lysis buffer. Polysomes were digested with 2.5 μg RNase A (Thermo Scientific, EN0531) per 750 ng total RNA for 20 min at 25°C at 750 rpm. The reaction was quenched with 0.4 mg/mL heparin. 3 mL of the lysate was layered on top of 9 mL 25% sucrose cushion in gradient buffer (25% sucrose, 20 mM HEPES pH 7.3, 150 mM KCl, 5 mM MgCl_2,_ 100 mg/mL cycloheximide, 1x EDTA-free protease inhibitor cocktail). Ribosomes were pelleted by centrifugation at 100,000 g overnight at 4°C in a SW40Ti rotor. After the supernatant was removed, the ribosomal pellet was resuspended with 600 μL of resuspension buffer (10% glycerol, 20 mM HEPES pH 7.3, 150 mM KCl, 5 mM MgCl_2,_ 0.1% NP-40, 100 mg/mL cycloheximide, 1x EDTA-free protease inhibitor cocktail).

For co-IP 100 μg of Monoclonal ANTI-FLAG® M2 antibody (Sigma, F1804) was coupled to the 400 μL of protein G Dynabeads (Invitrogen) in 1 mL of lysis buffer without DTT and cycloheximide for 20 min rotating at room temperature, washed three times with 1 mL of 25% sucrose in gradient buffer supplemented with 0.1% NP-40, resuspended in the original bead volume and added to 500 μL of resuspended ribosomal pellet. The IP was incubated on a rotator at +4°C for overnight, washed 4 times with wash buffer (10% glycerol, 20 mM HEPES pH 7.3, 150 mM KCl, 5 mM MgCl_2,_ 0.01% NP-40, 100 mg/mL cycloheximide, 1x EDTA-free protease inhibitor cocktail), followed by 5 washes with (20mM HEPES, KCl 150mM). The samples were eluted from beads using tryptic digestion and the peptides were submitted for mass spectrometry at Vienna BioCenter Proteomics Core Facility.

100 μL aliquot of ribosomal pellet sample was taken as an IP input control. It was filled with dH_2_O up to 1 mL, 10 μL of 10% sodium deoxycholate were added and vortexed. Proteins were precipitated on ice for 10 min after addition of 200 μL of 50% TCA and centrifuged at 20,000 g for 20 min at 4°C. Pellet was washed twice with 1 mL of 100% acetone, all supernatant was discarded and the pellet was vacuum-dried briefly for 5 min at 45°C vacuum evaporator. Samples were prepared and tryptic digested, using the iST kit (PO 00001, PreOmics) according to the manufacturer’s instructions. 250 ng of each peptide sample were submitted for mass spectrometry at Vienna BioCenter Proteomics Core Facility. For the IP analyses, we used a cut off of at least 10% coverage of a protein of interest.

### Mammalian siRNA and plasmid transfections

For reporter accumulation experiments, RKO iCas9 cells were seeded in 6-well plates and treated with 450 nM doxycycline for 48 h to induce expression of Cas9. The cells were then transfected with 2 μg plasmid using jetOPTIMUS® reagent according to the manufacturer’s instructions (Avantor, 101000051). Cells were washed with cold PBS buffer and collected on plate in RIPA buffer (150 mM NaCl, 1% NP-40, 0.5% Sodium deoxycholate, 0.1% SDS, and 25 mM TRIS pH 7.4) with 1x EDTA-free protease inhibitor cocktail (Roche, 11873580001). Cell lysate was clarified on a table-top centrifuge at maximum speed (20,000 g) for 20 min at 4°C.

To test the effect of RQC components’ loss on UFMylation, HCT116 tetON OsTIR1 cells were seeded in 24-well plates and transfected with Dharmacon ON-TARGETplus Human siRNA SMARTpools at 10 nM against ASCC3 (L-012757-01-0005) and at 25 nM against PELO (L-019068-01-0005) and ZNF598 (L-007104-00-0005). ON-TARGETplus Non-targeting Control Pool (D-001810-10-05) at 25 nM was used as a control. Cells were passed to 6-well plates and either transfected with 2 μg of plasmids containing CytoK0, CytoK20, ERK0, or ERK20 reporters ^28^ for 24 h using jetOPTIMUS® reagent (Avantor, 101000051) according to the manufacturer’s instructions or treated with 4 μM anisomycin dissolved in DMSO for 1 h before collection.

RKO iCas9 cells were seeded in 10-cm dishes in media supplemented with 450 nM doxycycline for 48h to induce expression of Cas9. Cells were then either transfected with 10 μg of plasmids containing CytoK0, CytoK20, ERK0, or ERK20 reporters ^28^ for 24 h using jetOPTIMUS® reagent according to the manufacturer’s instructions (Avantor, 101000051), or treated with 4μM anisomycin dissolved in DMSO for 1 h before collection.

### Flow cytometry analysis

HCT116 tetON OsTIR1 cells were seeded in 24-well plates and transfected with Dharmacon ON-TARGETplus Human siRNA SMARTpools for 72 h total as described in “Mammalian siRNA and plasmid transfections”. Cells were passed to 6-well plates and transfected with 2 μg of pEGFP-N1 plasmid containing CytoK0, CytoK20, ERK0, or ERK20 reporters ^28^ for 24 h using jetOPTIMUS® reagent according to the manufacturer’s instructions (Avantor, 101000051). Cells were harvested in trypsin, resuspended in full media and analysed by flow cytometry (BD LSRFortessa™ Cell Analyser). Data was analysed in FlowJo software.

### Cell lysis and immunoblotting

Clear cell lysates were collected in RIPA buffer with 1xEDTA-free protease inhibitor cocktail (Roche, 11873580001), clarified on a table-top centrifuge at maximum speed (20,000 g) for 20 min at 4°C, and analysed on 10%, 12% or 15% SDS-polyacrylamide gels, followed by a semi-dry transfer (BioRad blotter) or wet transfer (120V, 1 h 20 min on a BioRad blotter) on a 0.2 μm nitrocellulose membrane. Membrane was blocked in 1xTBS buffer with 0.1% Tween containing 5% milk for 1 h at room temperature. Primary antibody (listed in Supplementary table 2) was diluted in 2.5% milk 1x TBS-0.1% Tween buffer and incubated either 1 h at room temperature or at 4°C overnight, followed by three washes with 1x TBS-0.1% Tween buffer. Secondary antibody conjugated with horseradish peroxidase (Promega) was diluted in 2.5% milk 1x TBS-0.1% Tween buffer and incubated for 1 h at room temperature, followed by three washes with 2.5% milk 1x TBS-0.1% Tween buffer. The blots were developed using chemiluminescent detection on iBright ThermoFisher machine and quantified in iBright Analysis Software (ThermoFisher).

### Immunoprecipitations

For detecting UFL1 binding to nascent chain-containing 60S ribosomes, RKO iCas9 cell lines expressing gRNA against AAVS1 (non-targeting control), NEMF, or UFM1 (gRNA sequences are listed in Supplementary table 2) were seeded in 10-cm dishes (2 dishes per condition) in media supplemented with 450 nM doxycycline for 48h to induce expression of Cas9. Cells were then transfected with 10 μg of plasmids containing CytoK20, or ERK20 reporters ^28^ for 24 h using jetOPTIMUS® reagent according to the manufacturer’s instructions (Avantor, 101000051). Cells were then washed in cold PBS supplemented with 100 μg/mL cycloheximide, lysed on the plate with ice-cold lysis buffer (20 mM HEPES pH 7.3, 120 mM KCl, 5 mM MgCl_2_, 1% NP-40, 100 μg/mL cycloheximide, 0.5 mM DTT, 1x EDTA-free protease inhibitor cocktail). Cell lysates were further incubated on ice for 10 min with intermittent vortexing, passed three times through the 27G needle, and clarified on a table-top centrifuge at maximum speed (20,000 g) for 20 min at 4°C. 700 μL of the lysate was layered on top of 233 μL of sucrose cushion (1 M sucrose, 20 mM HEPES pH 7.3, 120 mM KCl, 5 mM MgCl_2,_ 100 μg/mL cycloheximide, 0.5 mM DTT). Ribosomes were pelleted by centrifugation at 100,000 g for 1,5 h at 4°C in a TLA100.3 rotor. After the supernatant was removed, the ribosomal pellet was resuspended with 150 μL of lysis buffer. Immunoprecipitations were performed with 40 μL of resuspended GFP-Trap® Magnetic Particles (ChromoTek, gtd-20) per condition, the mixture was incubated on a rotator at +4°C for 1 h, washed 3 times with wash buffer (20 mM HEPES pH 7.3, 195 mM KCl, 5 mM MgCl_2,_ 0.5% NP-40, 100 μg/mL cycloheximide, 0,1 mM DTT, 1x EDTA-free protease inhibitor cocktail), and eluted in 30 μL of 1xSDS sample buffer without DTT. Samples were incubated at 70°C for 10 min, the eluate was separated from the magnetic particles, DTT was added to the final concentration of 10 mM, samples were incubated at 95°C for 5 min and loaded on a SDS-PAGE gel followed by Western blotting.

### Plant experiments

All *Arabidopsis thaliana* lines used in this study originate from the Columbia (Col-0) ecotype. Mutant lines used in this study are listed in Supplementary table 2. Seedlings were grown in liquid 1/2 MS medium containing 1% sucrose under 16 h light/8 h dark photoperiod for 7 days with shaking at 80 rpm. 7-day-old seedling grown in liquid 1/2 MS medium were treated with 100 μM anisomycin (ANS) for 16 hours under continuous light with shaking at 80 rpm. An equal volume of pure dimethyl sulfoxide (DMSO) was added as control. Subsequently, seedlings were then frozen in liquid nitrogen after chemical treatment and homogenized for western blotting. The total protein was extracted with Grinding Buffer (50 mM Tris-HCl, 150 mM NaCl, 1 % Glycerol, 0.5 % NP-40, 1.5 mM MgCl_2_, 1x protease inhibitor cocktail). The SDS loading buffer was added to lysates, and the samples then were boiled at 95°C for 10 minutes. 10 μg of sample was loaded per lane. SDS-PAGE was performed using gradient 4– 20% Mini-PROTEAN TGX Precast Protein Gels (BioRad). Blotting on nitrocellulose membranes was performed using a semi-dry Turbo Transfer Blot System (BioRad). The membranes were blocked with 5 % skimmed milk in TBS and 0.1 % Tween 20 (TBS-T) for 1 hour at room temperature. Subsequently, the membranes were incubated with primary antibody, followed by incubation with secondary antibody conjugated to horseradish peroxidase (HRP). After three times 10 minutes washes with TBS-T, the immune-reaction was developed using ECL SuperSignal West Femto (Thermo) and detected with iBright Imaging System (Thermo). Protein bands intensity was quantified using the iBright Imaging analysis System (Thermo). The average relative intensities and a standard error were calculated from three biological replicates.

### Statistical analysis and reproducibility

All experiments were performed at least 3 times unless otherwise indicated in figure legends. Statistical analysis was performed in GraphPad Prism 10 software using Student t-test.

